# A Network Pharmacology Approach to Elucidate the Anti-inflammatory Effects of Ellagic Acid

**DOI:** 10.1101/2023.01.10.523505

**Authors:** Skyler H. Hoang, Hue Dao, Emerson My Lam

## Abstract

Ellagic acid (EA) is a naturally occurring polyphenolic compound found in various fruits and vegetables like strawberries, raspberries, pomegranates, and nuts such as pecans and walnuts. With its antioxidant properties, EA has shown potential health benefits, although further research is necessary to fully comprehend its effects, mechanisms, and safe and effective application as a complementary medicine. Notably, there is accumulating evidence of EA’s anti-inflammatory effects; however, the precise underlying mechanism remains unclear. To investigate the anti-inflammatory properties of EA, a network pharmacology approach was employed. The study identified 52 inflammation-related targets of EA and revealed significant signaling pathways and relevant diseases associated with inflammation through GO and KEGG analysis. Furthermore, topological analysis identified 10 important targets, including AKT1, VEGFA, TNF, MAPK3, ALB, SELP, MMP9, MMP2, PTGS2, and ICAM1. Molecular docking and molecular dynamics simulations (integrated with were conducted molecular mechanics Poisson-Boltzmann), indicating that AKT1, PTGS2, VEGFA, and MAPK3 are the most likely targets of EA. In summary, this study not only confirmed the anti-inflammatory effects of EA observed in previous research but also identified the most probable targets of EA.

## Introduction

Ellagic acid (EA) is a naturally occurring polyphenolic compound with antioxidant properties found in a variety of fruits and vegetables, including strawberries, raspberries, pomegranates, and nuts such as pecans and walnuts (Baliga et al., 2019; Sharifi-Rad et al., 2022). EA has been studied for its potential health benefits, including its ability to support the immune system and protect cells from damage (Sharifi-Rad et al., 2022). Some research has suggested that EA may have anti-inflammatory and chemopreventive effects, which could potentially help to reduce the risk of chronic diseases such as heart disease and cancer (BenSaad et al., 2017; Päivärinta et al., 2006; Ridker, 2017). For instance, in a study model of culture of primary human gingival epithelial cells, EA was demonstrated to regulate the oral innate immunity via increasing the expression of chemokine ligand 5, IL-2, IL-1β and decreasing the expression of IL-6, IL-8, and TNF (Promsong et al., 2015). In terms of its potential anticancer effects, some research has suggested that ellagic acid may help to inhibit the growth of cancer cells and induce cell death in certain types of cancer, such as breast, colon, and prostate cancer (Ceci et al., 2018). In another instance, transcriptional profiling using microarray and functional analysis on Caco-2 cells exposed to EA revealed that EA could affect the expression of growth factor receptors (fibroblast growth factor receptor 2, epidermal growth factor receptor), oncogenes (K-Ras, c-Myc), and tumor suppressors (dual specificity phosphatase 6, Fos) and of genes involved in cell cycle (cyclin B1, cyclin B1 Interacting Protein 1) (González-Sarrías et al., 2009). EA is also claimed to support a variety of other health goals, including weight loss, cardiovascular health, and improved digestion (Kang et al., 2016). More research into the mechanism of this compound is needed to fully understand these potential effects and how anti-inflammatory, anti-cancer, or metabolic syndrome preventive effects of EA can be utilized. Overall, EA appears to have potential health benefits, but more research is needed to fully understand its effects and how it can be used safely and effectively as a complimentary medicine.

On another note, inflammation is a natural process that occurs in the body in response to injury or infection (L. Chen et al., 2018). It is characterized by swelling, redness, warmth, and pain, and is a crucial part of the body’s immune response (Scott, 2004). When the body is injured or infected, immune cells are activated to help fight off the infection or repair the damage (Fang et al., 2018; Luster et al., 2005). This process involves the release of various chemicals and signaling molecules that cause inflammation in the affected area. Inflammation is important because it helps to protect the body from harm and promote healing. However, chronic inflammation, which is a long-term, persistent form of inflammation, can be harmful and contribute to the development of chronic diseases such as heart disease, diabetes, and cancer (Furman et al., 2019). Chronic inflammation is often caused by underlying conditions such as obesity, smoking, and high levels of stress, and can also be triggered by certain foods and environmental factors (Bosma-den Boer et al., 2012). There is increasing evidence to suggest that chronic inflammation is a key factor in the development of many chronic diseases. For instance, cardiovascular diseases are often caused by inflammation in the arteries, which can lead to the build-up of plaque and eventually cause a heart attack or stroke (Kabbany et al., 2016).

Similarly, inflammation in the pancreas may contribute to the development of diabetes, and inflammation in the colon may increase the risk of colon cancer (Terzić et al., 2010). Studying inflammation is important because it helps to understand the underlying causes of chronic diseases and may lead to the development of new treatments and prevention strategies. Certain medications, such as nonsteroidal anti-inflammatory drugs and corticosteroids, can help to reduce inflammation and pain or may be used to treat conditions such as arthritis (Del Grossi Moura et al., 2018). However, their misusage has been associated with adverse events such as gastrointestinal ulcers, serious cardiovascular events, or hypertension (Vonkeman & van de Laar, 2010; Yasir et al., 2022). It should be noted that lifestyle changes such as eating a healthy diet, exercising regularly, and managing stress may also help to reduce inflammation and lower the risk of chronic diseases (Margină et al., 2020). EA, with accumulating evidence of anti-inflammatory properties, can be easily found as a part of a healthy, plant-based diet (Kang et al., 2016). This investigation is also further compelled by the fact that EA has been demonstrated to exert glucose-lowering, antioxidant, anti-inflammatory, and anti-glycation effects in murine diabetic models (Amor et al., 2020). Herein, a network pharmacology approach, integrated with molecular docking and molecular dynamics simulation, is utilized to elucidate the mechanisms underlying anti-inflammatory properties of EA.

## Results and Discussion

### 1. Network Construction and Analysis

The retrieval of EA target proteins and target proteins for inflammation results in 157 and 886 target proteins, respectively (**Figure 1**). There were 52 overlapped proteins, representing potential therapeutic targets of EA for its anti-inflammatory properties. A PPI network of 52 overlapped was constructed and resulting in 325 number of edges, 12.5 average node degree, 0.688 average local clustering coefficient, as well as a PPI enrichment p-value of less than 1.0e-16, indicating that the proteins are biologically connected (**Figure 1**).

**Figure 1.**
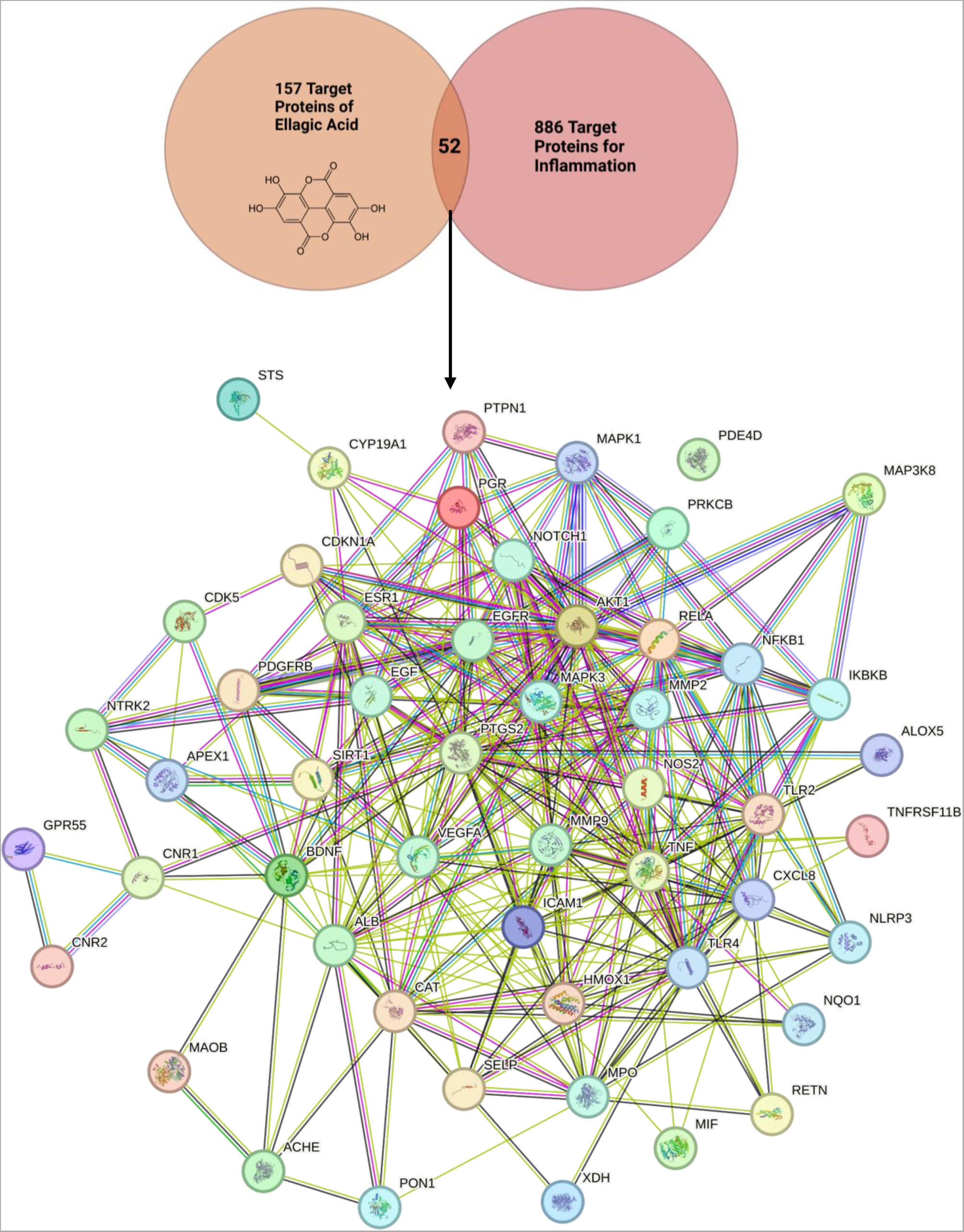
Venn diagram showing the overlap between 157 target proteins of ellagic acid (EA) and 886 target proteins for inflammation as well as the protein-protein interaction network of 52 overlap proteins via string-db.org.

The 5 most significant GO Biological Process enriched pathways include: response to organic substance, response to oxygen-containing compound, response to lipid, response to chemical, and cellular response to chemical (**Figure 2A**). The 5 most significant GO Molecular Function enriched pathways include: identical protein binding, protein binding, signaling receptor binding, enzyme binding, and ion binding (**Figure 2A**). The 5 most significant GO Cellular Component enriched pathways include: vesicle lumen, endomembrane system, secretory granule, secretory granule lumen, and membrane raft (**Figure 2A**).

**Figure 2.**
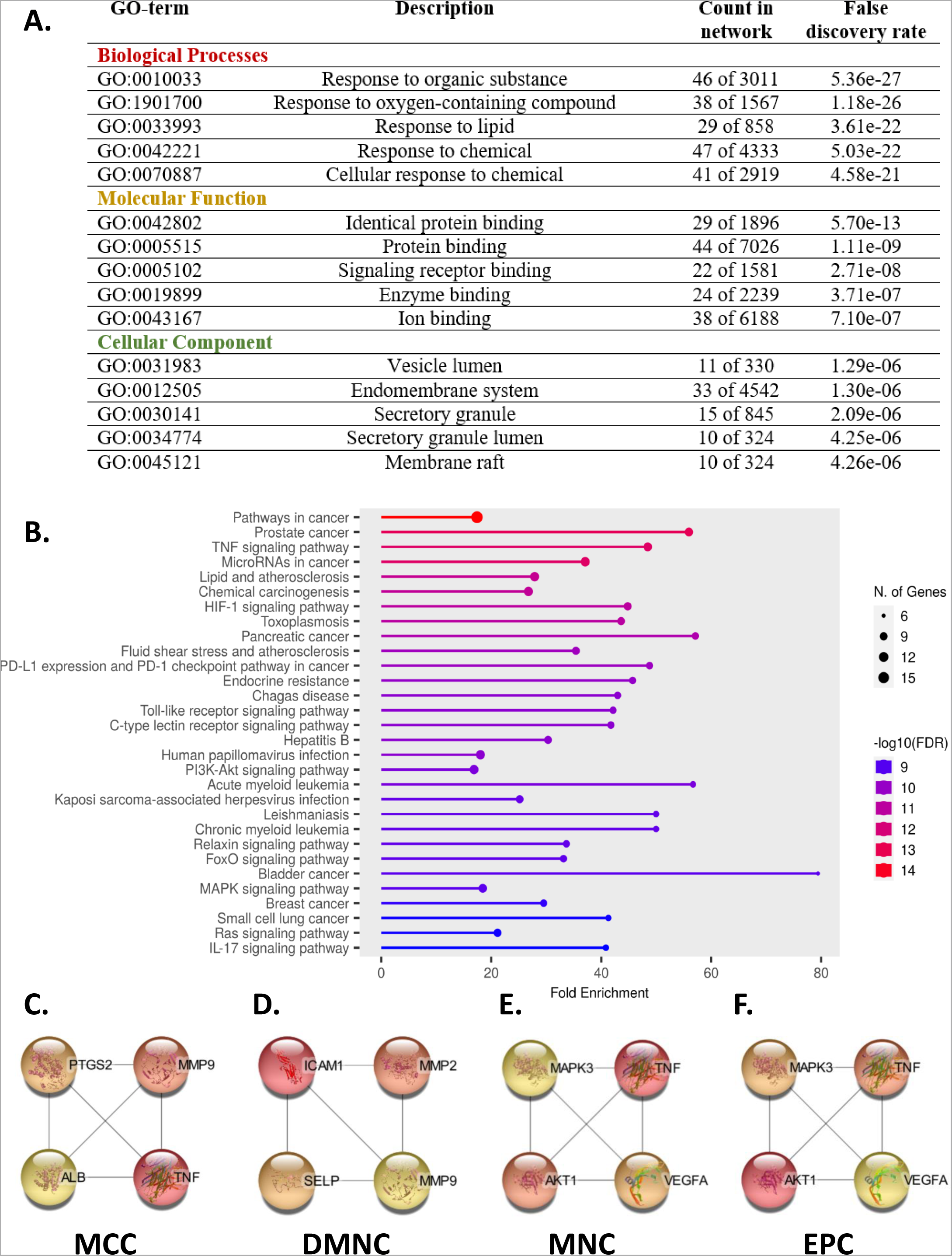
Enrichment analysis and topological analysis of the protein-protein interaction. network. (A) Gene Ontology (GO) enrichment analysis. (B) Kyoto Encyclopedia of Genes and Genomes (KEGG) enrichment analysis. Topological analysis to determine important proteins of the network via (C) Maximal Clique Centrality (MCC) (D) Density of Maximum Neighborhood Component (DMNC) (E) Maximum Neighborhood Component (MNC) (F) Edge Percolated Component (EPC)

Interestingly, lipid regulation and inflammation have an intimate relationship. Particularly, abnormalities in lipid profile or regulation can lead to abnormalities in inflammatory responses, and the reverse is also true in that alterations of inflammatory signaling (in the context of metabolic diseases such as diabetes or atherosclerosis) can result in changes of lipid metabolism in liver, skeletal muscle, macrophage, or adipose tissue (Glass & Olefsky, 2012). In a murine basal diet model, EA supplementation decreased total cholesterol, low-density lipoprotein, and the abundance of acetyl-CoA carboxylase while increasing high-density lipoprotein and the abundances of phospho-hormone-sensitive lipase, carnitine palmitoyl transferase 1B, and peroxisome proliferator-activated receptor alpha protein (Xu et al., 2021). In a randomized double-blind clinical trial, patients with type 2 diabetes consuming 180 mg of EA daily for 8 weeks demonstrated a decrease in inflammatory markers (i.e, TNF, and IL-6), total cholesterol, triglycerides, as well as low-density lipoprotein and an increase in total antioxidant capacity, glutathione peroxidase activity, as well as superoxide dismutase enzymes (Ghadimi et al., 2021). ROS are crucial signaling molecules that play a vital role in the course of inflammatory diseases. EA effectively reduces endothelial ROS levels and improves arterial relaxation impairment caused by high glucose in a model of intact rat aortas and human aortic endothelial cells activated with high glucose concentration (Rozentsvit et al., 2017). A wide body of literature demonstrated that polyphenolic compounds can be helpful as adjuvant therapy due to their potential anti-inflammatory effects, antioxidant activity, and inhibition of enzymes involved in eicosanoids formation (Hussain et al., 2016). It is within reasons to speculate that EA might counteract inflammation and tissue damage via this pathway, especially when more recent evidence also suggests that EA, in a model of zebrafish embryonic development, might provide protective effects against DNA oxidative damage while simultaneously increasing the embryo survival rate and improving morphological parameters of the larvae (Mottola et al., 2020).

Interestingly, the vesicle lumen was identified by GO Cellular Compartment analysis as a potential cellular compartment where all these processes might take place. Aforementioned results of previous studies are highly related to cellular production and release of inflammatory mediators, a critical component of vesicle lumen (Türk et al., 2010). Previous network pharmacology utilizing bioinformatics also predicted vesicle lumens as potential target of EA in the model of endometriosis (Wu et al., 2022) and beta amyloid 25-35 peptide-induced neurotoxicity (Li et al., 2022). Interestingly, EA was also predicted by GO Cellular Function to potentially be involved in ion binding process. Within the scope of inflammation, H+/K+ ATPase activity, known to be stimulated by inflammatory and other factors and by the bacterium *Helicobacter pylori* (Z. Zhang et al., 2020), was demonstrated to be inhibited by EA hog gastric with a 50% inhibition at 2.1 × 10å-6 M (Murakami et al., 1991), postulating a compelling case to study the intersection of EA, inflammatory mediators, vesicle lumens, and ion exchange process.

The 30 most significant KEGG enriched pathways are illustrated in **Figure 2B**. Notably, cancer-related pathways include: pathways in cancer, prostate cancer, microRNAs in cancer, chemical carcinogenesis, pancreatic cancer, acute myeloid leukemia, chronic myeloid leukemia, bladder cancer, breast cancer, and small cell lung cancer (**Figure 2B**). Cancer and inflammation have a uniquely inseparable relationship: cancer growth and progression are strongly associated with inflammation, which, in turn, can affect cancer formation, metastasis or response to therapy (Singh et al., 2019; Zhao et al., 2021). Recent evidence from *in vitro* and *in vivo* studies have suggested that EA may be able to inhibit tumor cell proliferation, induce apoptosis, break DNA binding to carcinogens, block virus infection, and disturb inflammation, angiogenesis, and drug-resistance processes, which are all important factors in the development and progression of cancer (H.-M. Zhang et al., 2014). Other pathways relevant to inflammation also include Hepatitis B, FoxO, TNF, and HIF-1 signaling pathways. In conjunction, it was demonstrated that EA could effectively inhibit Hepatitis B-infected HepG2 2.2.15 cells from secreting hepatitis B e-antigen (Shin et al., 2005). In a model of high glucose-induced injury in rat mesangial cells, EA was demonstrated to inhibit the activation of the PI3K/Akt signaling pathway and reduce the expression levels of downstream transcription factor FOXO3a (W. Lin et al., 2021). In a model of 1,2-dimethylhydrazine-induced rat colon carcinogenesis, EA downregulated iNOS, PTGS2, TNF and IL6 via inhibition of NF-kB and demonstrated chemopreventive effects on colon carcinogenesis (Umesalma & Sudhandiran, 2010). Lastly, in a study using *in vitro* model of lung cancer (i.e., HOP62 and H1975 cell lines) EA decreased both mRNA expression of target genes of HIF-1 as well as the protein level of HIF-1, whose inhibition may enhance the effectiveness of a vast variety of innovative and established anticancer medicines (Duan et al., 2020).

To further determine critical proteins in EA’s anti-inflammatory activity, topological analysis was performed (**Figure 2C-F**) using cytoHubba MCC, DMNC, MNC, and EPC algorithms. The MCC algorithm determined PTGS2, MMP9, ALB, and TNF as the 4 most essential proteins of the network (**Figure 2C**). Alternatively, the DMNC algorithm determined ICAM1, MMP2, SELP, and MMP9 as the 4 most essential proteins of the network (**Figure 2D**). Furthermore, the MNC algorithm determined MAPK3, TNF, AKT1, and VEGFA as the 4 most essential proteins of the network (**Figure 2E**). Lastly, the EPC algorithm determined MAPK3, TNF, AKT1, and VEGFA as the 4 most essential proteins of the network (**Figure 2F**). In essence, per the recommendation of Chin et al., 2014, combining the results of these 4 algorithms determined that the top 10 most important proteins of the network are PTGS2, MMP9, MMP2, ALB, TNF, ICAM1, SELP, MAPK3, VEGFA, and AKT1. PTGS2 is an enzyme that plays a role in bio-transforming arachidonic acid to prostaglandins, which are signaling molecules that have various functions in the body, including the regulation of inflammation (Simon, 1999). MMP9, in the context of inflammation, is often upregulated and released by both immune cells (i.e, neutrophils and monocytes) as well as endothelial or fibroblast cells (Al-Sadi et al., 2021; Yabluchanskiy et al., 2013). Notably, MMP9 was demonstrated maintain epithelial barrier function and preserve mucosal lining in colitis associated cancer in a model of murine colonic epithelium and CaCo2BBE intestinal cell line (Pujada et al., 2017). The upregulation and release of MMP9 can eventually contribute to the breakdown of extracellular matrix proteins and other proinflammatory molecules, leading to tissue damage. In a similar manner, MMP2 is a protease that can cleave several extracellular matrix proteins, including collagens, laminins, and elastins and is important for various physiological processes, such as tissue repair and remodeling, as well as pathological processes such as cancer cell invasion and metastasis (Maybee et al., 2022; Song et al., 2021). Inadequacy of ALB is associated with inflammatory conditions (Soeters et al., 2019); a clinical study demonstrated that ALB is negatively correlated with inflammatory indices (i.e, C-reactive protein and white blood cell count), implying a potential role of ALB in the regulation of inflammatory and immune responses (Sheinenzon et al., 2021). TNF mediates inflammation by activating immune cells and inducing the expression of other cytokines, chemokines, and adhesion molecules and eventually clears infections and in the process of tissue repair after injury, but TNF can also be the driving force of chronic inflammation, leading to tissue damage and the development of autoimmune diseases (Chu, 2013; Kalliolias & Ivashkiv, 2016). Within the same vein, EA reversed the release of IL-6 and IL-8 induced by TNF in Caco-2 cell (Iglesias et al., 2020). ICAM1 is a type of cell adhesion molecule that is expressed on the surface of various types of cells, including epithelial cells, endothelial cells, and immune cells and plays a critical role in the inflammatory response by mediating the adhesion of leukocytes to the endothelium (Bui et al., 2020). SELP is a type of cell adhesion molecule that is expressed on the surface of stimulated endothelial cells and activated platelets; SELP can mediate the interaction between leukocytes and endothelial cells, allowing leukocytes to “roll” along the surface of the endothelium, which is important for the recruitment of leukocytes to sites of inflammation (M. Chen & Geng, 2006). MAPK3 plays a key role in regulating various cellular processes, including cell proliferation, differentiation, and survival and has been shown to be upregulated in immune cells during inflammation, and its activation also is associated with the production of pro-inflammatory mediators, such as cytokines and chemokines (Lucas et al., 2022). In inflammation, VEGFA is important for recruiting of immune cells to the site of inflammation, forming of new blood vessels, maintaining the integrity of blood vessels, wound healing, and the development of the placenta during pregnancy; however, excessive, or uncontrolled production of VEGFA can contribute to the development of certain diseases, such as cancer (Scaldaferri et al., 2009; Shaik-Dasthagirisaheb et al., 2013). Lastly, activation of AKT1 is known to lead to the activation of NF-κB, which in turn upregulates and encodes cytokines, chemokines, and adhesion molecules that play critical roles in inflammatory responses (Luthra et al., 2008; Zhou et al., 2013). EA might very well directly interact with these targets to exert its anti-inflammatory effects, as previous studies demonstrated that EA could affect the gene as well as protein expression of these targets (Attilio et al., 2010; Kowshik et al., 2014; C. Lin et al., 2019; Pattanayak et al., 2017; Umesalma & Sudhandiran, 2010; Yu et al., 2007). Therefore, molecular docking and molecular dynamics simulations were utilized to validate said speculation.

### 2. Molecular Docking and Molecular Dynamics Simulation Studies

Results of molecular docking (binding affinity, predicted Ki, and contact surface) are summarized in **Table 1**. In brief, from the molecular docking results, EA had the strongest binding affinities for AKT1, MAPK3, PTGS2 (with –9.894, –9.452, and –9.359 kcal/mol, respectively) and weakest binding affinities for ICAM1, TNF, and VEGFA (with –6.871, –6.479, –6.160 kcal/mol, respectively) (**Table 1**).However, the crucial analyses of this study rely on the ligand contact analysis (**Figure 3**) and MM/PBSA-calculated binding energies (**Figure 4A**) by means of molecular dynamics simulations. EA was able to maintain continuous contacts with all important target protein throughout 100 ns of molecular dynamics simulations (**Figure 3**). From MM-PBSA binding energy calculations, EA demonstrated strongest binding affinities with AKT1, PTGS2, VEGFA, MAPK3 with –82.37 ± 159.8, –52.06 ± 171.4, –19.24 ±79.02, and – 13.98 ± 149.6 kJ/mol, respectively (**Figure 4A**). EA demonstrated weaker binding with ALB, MMP9, ICAM1, TNF, MMP2, and SELP with –9.597 ± 223.9, 16.84 ± 169.2, 29.11 ± 169.2, 46.92 ± 80.85, 54.92 ± 81.22, and 59.51 ± 92.92 kJ/mol, respectively (**Figure 4A**). However, these results do not necessarily negate ALB, MMP9, ICAM1, TNF, MMP2, and SELP as potential targets of EA, as it only highlights that AKT1, PTGS2, VEGFA, and MAPK3 are the most likely targets of EA. However, further analyses for the complexes with the highest MM/PBSA binding energies somewhat corroborate this result (**Figure 4B**). EA showed binding energies towards MAP3K and VEGFA that are comparable to known binders, except AKT1 and PTGS2. Specifically, the binding energies of 10DEBC (for AK± 171.2 andT1), resveratrol (for MAP3K), ibuprofen (for PTGS2), and sorafenib (for VEGFA) are –265.2 ± 161.2, –24.12 ± 145.2, –215.1 ± 171.2, and –41.76 ± 81.92. However, this does not necessarily suggest thar EA is a non-binder for AKT1 or PTGS2, as it could potentially act a mild modulator, providing complementary or preventative effects. Further molecular dynamics simulation analyses are provided as Figure S1-S5.

**Figure 3.**
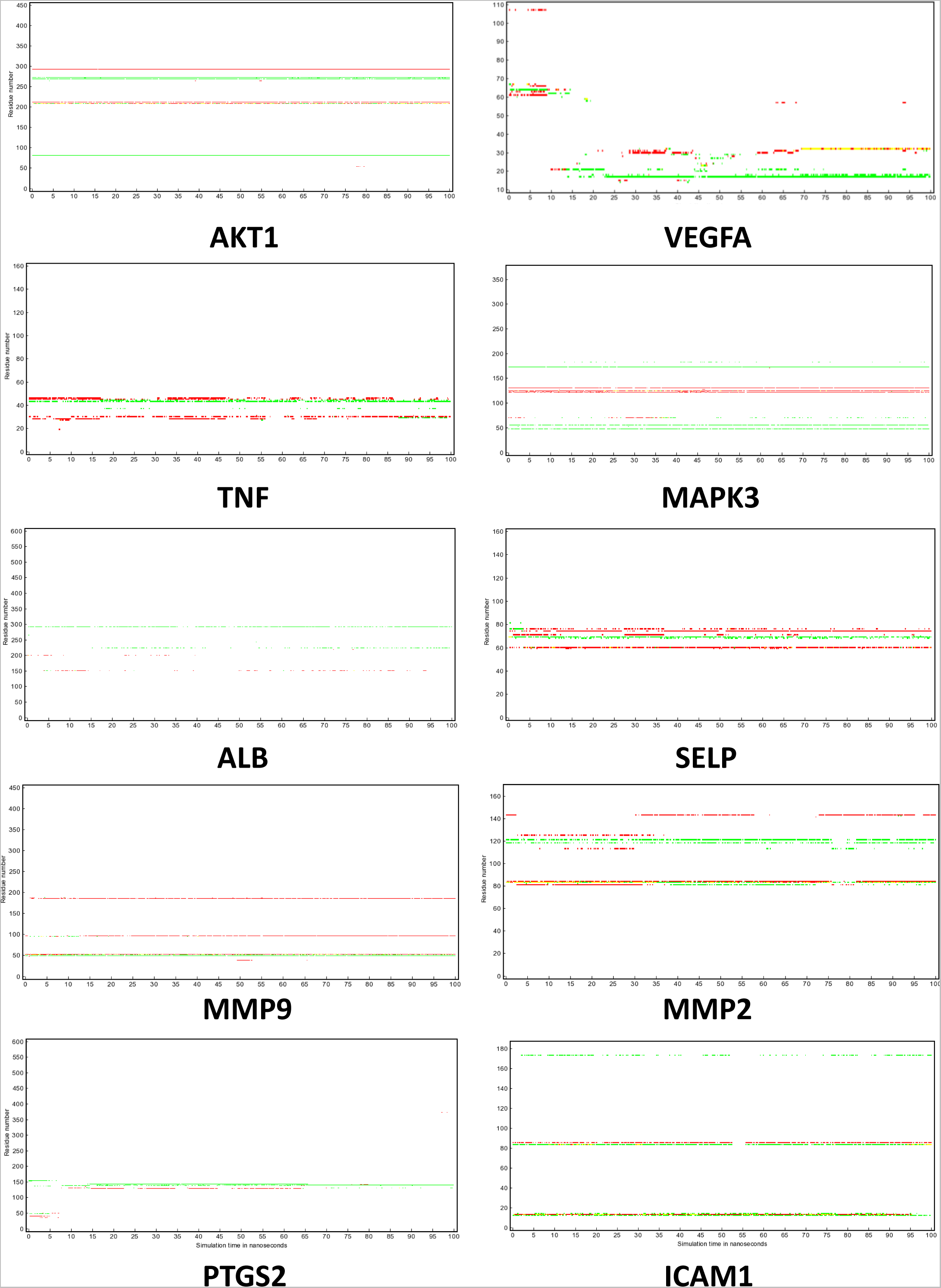
Residue contacts between ellagic acid (EA) and important proteins of the network determined through topological analysis demonstrating that EA can interact with putative target 100% of the time of the simulations. Red, green, and yellow colors denote hydrophobic interaction, hydrogen bonding, and the simultaneous combination of hydrophobic interaction and hydrogen bonding, respectively.

**Figure 4.**
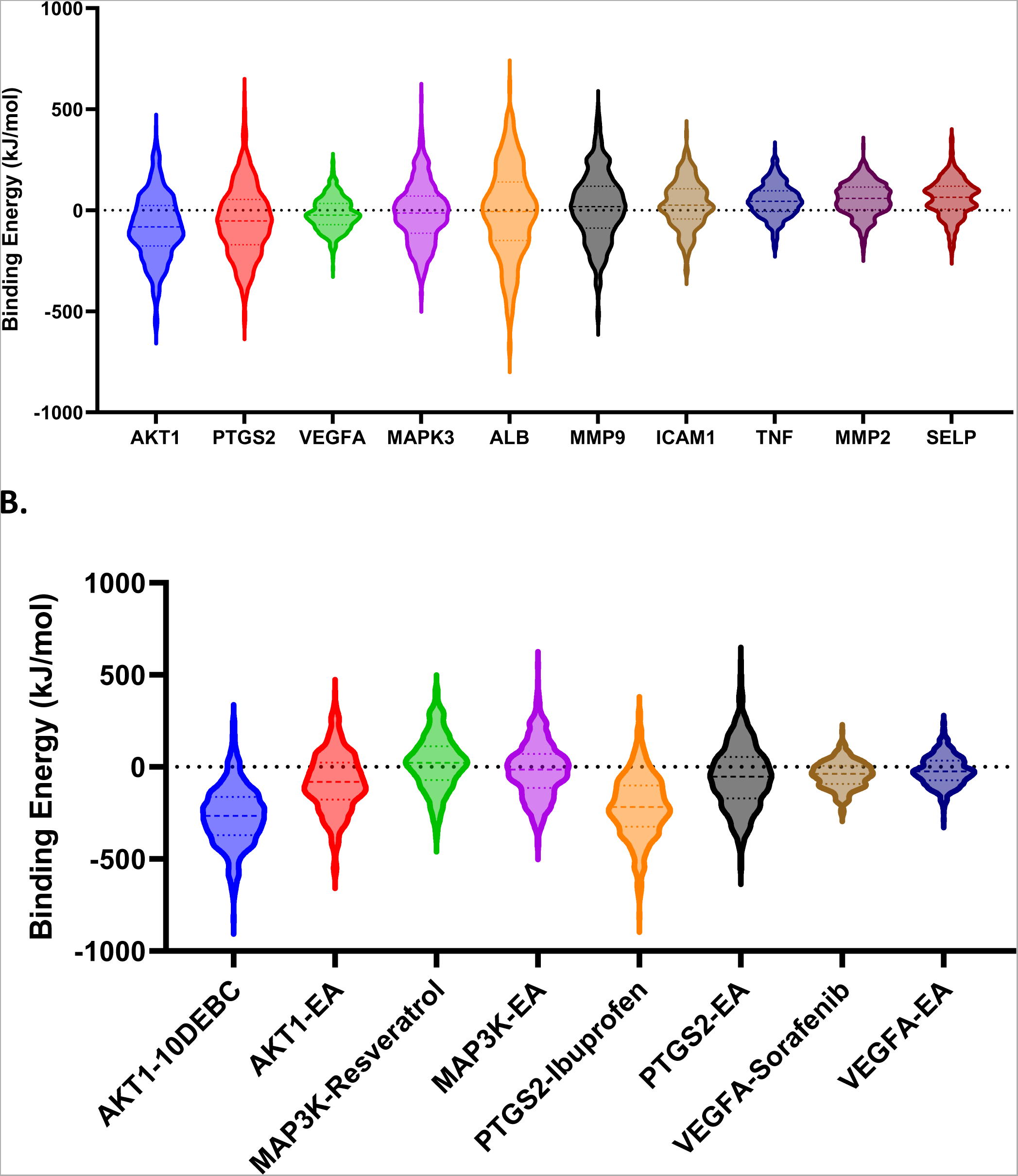
(A) Binding energies calculated by molecular mechanics Poisson-Boltzmann surface area (MM-PBSA) method, represented in violin plots in comparison with EA-protein complexes with the highest molecular docking binding energies. (B) Binding energies calculated by molecular mechanics Poisson-Boltzmann surface area (MM-PBSA) method, represented in violin plots in comparison with known binders.

**Table 1.**
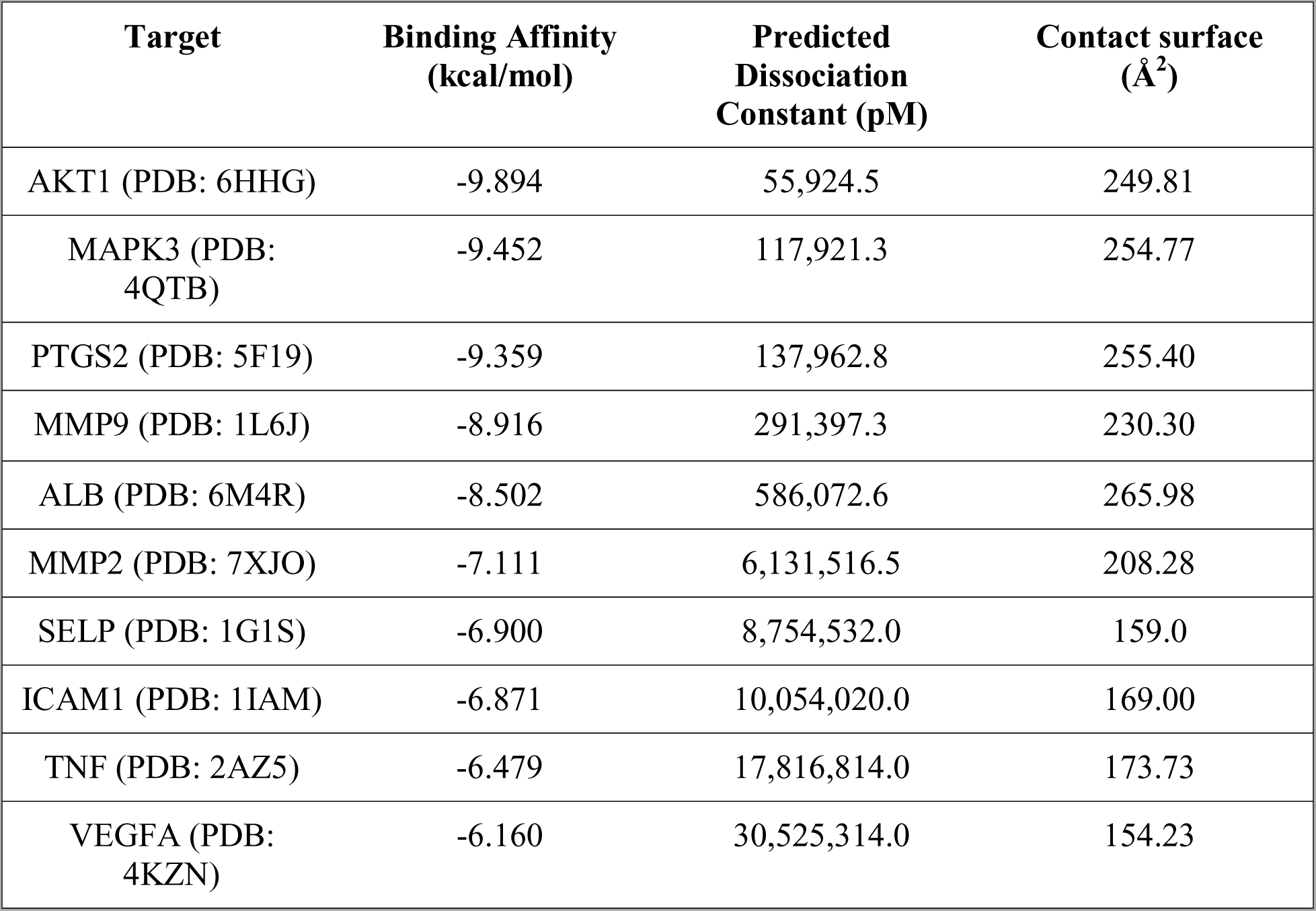
Results of molecular docking studies, sorted by binding affinities from strongest to weakest.

| Target | Binding Affinity<br>(kcal/mol) | Predicted<br>Dissociation<br>Constant (pM) | Contact surface<br>(Å<sup>2</sup>) |
| --- | --- | --- | --- |
| AKT1 (PDB: 6HHG) | -9.894 | 55,924.5 | 249.81 |
| MAPK3 (PDB:<br>4QTB) | -9.452 | 117,921.3 | 254.77 |
| PTGS2 (PDB: 5F19) | -9.359 | 137,962.8 | 255.40 |
| MMP9 (PDB: 1L6J) | -8.916 | 291,397.3 | 230.30 |
| ALB (PDB: 6M4R) | -8.502 | 586,072.6 | 265.98 |
| MMP2 (PDB: 7XJO) | -7.111 | 6,131,516.5 | 208.28 |
| SELP (PDB: 1G1S) | -6.900 | 8,754,532.0 | 159.0 |
| ICAM1 (PDB: 1IAM) | -6.871 | 10,054,020.0 | 169.00 |
| TNF (PDB: 2AZ5) | -6.479 | 17,816,814.0 | 173.73 |
| VEGFA (PDB:<br>4KZN) | -6.160 | 30,525,314.0 | 154.23 |

To reiterate, AKT1, PTGS2, VEGFA, MAPK3 had been demonstrated to play critical roles in progression of cancer, inflammation, HIF-1 signaling, and TNF signaling (Faurschou & Gniadecki, 2008; Fojtík et al., 2021; Guadagni et al., 2007; Lee et al., 2010; C.-C. Lin et al., 2004; Ramakrishnan et al., 2014; Siddappa et al., 2015; Stegeman et al., 2016). Thus, they, through topological analysis and molecular dynamics simulations, are most likely to be targets of EA for its anti-inflammatory activities and are included in the illustration of putative anti-inflammatory mechanism of EA (**Figure 5**). Particularly, TNF-alpha, regulated by NF-kB and responsible for the progression of cancer (Guadagni et al., 2007), was demonstrated by to stimulate PI3K-Akt in keratinocyte cell lines HaCaT and A431 cell lines, eventually resulting in enhanced mutagenesis and tumor development (Faurschou & Gniadecki, 2008). Compellingly, ellagic acid was demonstrated to provide protective effects against high glucose-induced injury in in rat mesangial cells via the PI3K/Akt/FOXO3a signaling pathway (W. Lin et al., 2021), which again further supports the findings of this study with predictive nature. Hypoxia and mild hypoxia cellular environments were demonstrated to increase PI3K-Akt and MAPK signaling and decrease PI3K-Akt signaling (respectively) in human pluripotent stem cells, demonstrating a unique but intimate relationship between hypoxia and PI3K-Akt and MAPK signaling (Fojtík et al., 2021). Within the same vein, in in vivo experiments using human esophageal epithelial cells (EPC2-hTERT), hypoxia conditions also upregulate PTGS2, but hypoxia conditions with cotreatment of IL-1β prevented said upregulation, demonstrating an intimate relationship between hypoxia, PTGS2, and inflammation. HIF-1 signaling pathway is also known for being the upstream regulator of VEGF (which is a critical biomarker for infection, cancer, and cardiovascular diseases). Furthermore, hypoxic conditions also induced AKT1 in neck squamous cell carcinoma and non-small-cell lung cancer cell lines (Stegeman et al., 2016). The nuance is also further extended, as it was previously demonstrated that MAP3K can potentially regulate ovulation by targeting PTGS2 (also a target of HIF-1 signaling pathway) in murine ovulating follicles (Siddappa et al., 2015). In short, we believe that EA might play a critical role in attenuating multiple “ends” or “streams” of these interconnected relationships. However, due to its, low hydrolysis and lipophilic and hydrophobic nature, EA is not readily absorbed in circulation but rather concentrated in the intestine (Mohammadinejad et al., 2022). We believe the intestine would be a great target for the investigation of animal or human models for the effects of EA dietary consumption on inflammation or diseases with inflammatory signatures.

**Figure 5.**
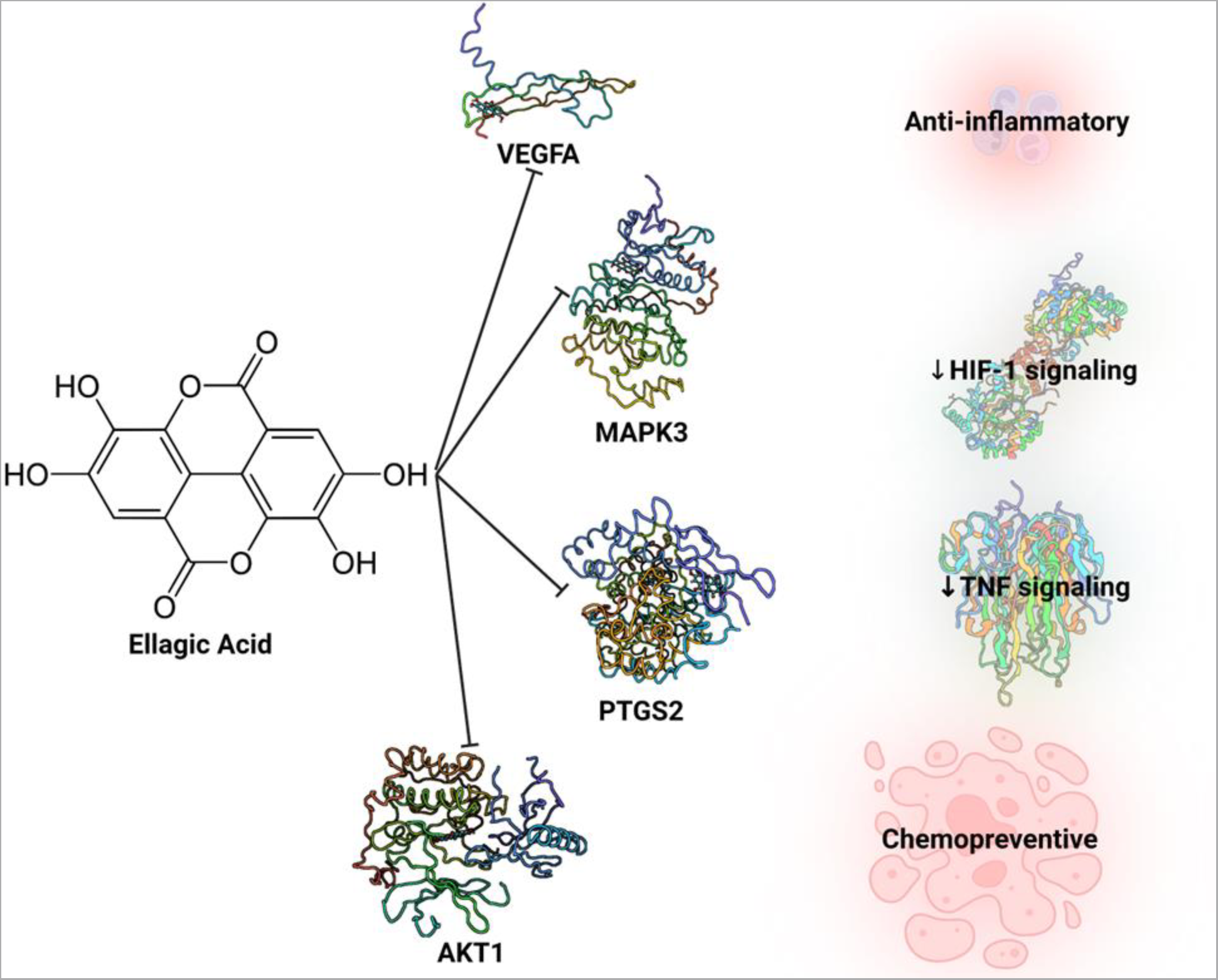
Illustrations of EA affecting inflammation, HIF-1 signaling, TNF signaling, and cancer pathways.

## Conclusion

In summary, a network pharmacology approach was used to understand the anti-inflammatory effects of EA. It was revealed that there are 52 targets of EA that are related to inflammation. Topological analysis determined 10 important targets that are AKT1, VEGFA, TNF, MAPK3, ALB, SELP, MMP9, MMP2, PTGS2, and ICAM1. Molecular docking and molecular dynamics simulations helped determine AKT1, PTGS2, VEGFA, MAPK3 as the most probable targets of EA. Computational studies need further experimental validations; however, this study corroborates with previous studies and provides potential targets and mechanisms for future research.

## Methods

### 1. Retrieval of EA target proteins

Target proteins of EA were retrieved from SwissTargetPrediction (Daina et al., 2019) and SymMap v2 (Wu et al., 2019) in UniProt ID format. String-db.org (Szklarczyk et al., 2021) was used to exclude any duplicate and resulted in a total of 157 EA target proteins.

### 2. Retrieval of target proteins for inflammation

DisGeNET (Piñero et al., 2019) was used to retrieve target inflammation proteins in UniProt ID format via the following accession numbers: C0021368 (“inflammation”), C0234251 (“inflammatory pain”), and C1290884 (“inflammatory disorder”). String-db.org was also used to exclude any duplicate and resulted in a total of 886 target proteins for inflammation.

### 3. Constructing PPI network of EA target proteins for anti-inflammatory properties

After two networks of 157 EA target proteins and 886 target proteins for inflammation were constructed on string-db.org, they were then sent to Cytoscape 3.9.1 (Shannon et al., 2003) to find mutual proteins between the two networks via the Merge-Intersection function, resulting in 52 EA target proteins that are also relevant for its anti-inflammatory properties. This visualization of this network and its basic topological analysis were then provided through string-db.org at a moderate confidence score of 0.55. Within string-db.org, GO enrichment analysis was also retrieved; 5 selected enriched terms with the lowest false discovery rate for each GO enrichment category (Biological Processes, Molecular Function, Cellular Component) were presented. On the other hand, ShinyGO 0.76.3 (Ge et al., 2020) was used to perform and visualize KEGG enrichment analysis.

### 4. Topological analysis of the PPI

Both local-based (i.e, MCC, DMNC, and MNC) and global-based (i.e, EPC) algorithms from the cytoHubba plug-in of Cytoscape were used to determine the most critical target proteins. The mathematical basis for these algorithms can be found at Chin et al., 2014. A protein is deemed significant if it ranks among the top four with the highest score in at least one category (i.e, MCC, DMNC, MNC, and EPC).

### 5. Molecular Docking

YASARA Structure (Land & Humble, 2018) was used to perform both molecular docking and molecular dynamics simulations. The dock_run.mcr script, which utilized Vina algorithm, was employed to perform molecular docking for EA towards important protein targets for inflammation. For each protein target, 100 docking runs were performed, and the docked pose with the most negative binding affinity (ΔG) in kcal/mol was selected for further validation through molecular dynamics simulations.

### 6. Molecular Dynamics Simulations

The md_run.mcr script was employed to perform molecular dynamics simulations for docked complexes to investigate the interactions between EA and its critical target proteins. In brief, the important parameters for the simulations are as follows: pH is 7.4; ion concentration is 0.9% NaCl; temperature is 298 K; water density is 0.997 g/ml; pressure control mode is solvent probe; cell shape is cube with periodic boundary; force field is AMBER14. The simulation is speed is ‘normal,’ in which a mixed-multiple timestep (of 2*1.25 femtoseconds) algorithm combined with LINCS algorithm is utilized. Eventually, analyses of contacts between EA and target proteins as well as movement RMSD of EA were retrieved through the md_analyze.mcr script. Binding energies derived from MM/PBSA were calculated using md_analyzebindenergy.mcr script, based on the following premise:

ΔG-binding = ΔG-complex – (ΔG-protein + ΔG-ligand) (in kJ/mol)

In which, ΔG-complex is calculated based on the solvation and potential energies of the complex; ΔG-protein is calculated based on the solvation and potential energies of the protein, and ΔG-ligand is calculated based on the solvation and potential energies of the ligand. More negative energies indicate better binding; positive energies do not indicate no binding. Top 4 complex with the highest MM/PBSA binding are further compared with known binders for validation.

## Data Availability

All predictive and enrichment data can be found in the Supplementary Excel File online. Sheet 1 corresponds to DisGeNet C0021368 (“inflammation”). Sheet 2 corresponds to C0234251 (“inflammatory pain”). Sheet 3 corresponds to C1290884 (“inflammatory disorder”). Sheet 4 corresponds to SymMap target of EA. Sheet 5 corresponds to SwissTargetPrediction target of EA. Sheet 6 corresponds to 52 targets of EA for inflammation. Sheet 7 corresponds to GO Biological Process enrichment analysis. Sheet 8 corresponds to GO Cellular Component enrichment analysis. Sheet 9 corresponds to GO Molecular Function enrichment analysis. Sheet 10 corresponds to KEGG enrichment analysis.

## Figure Captions

**Figure S1.**
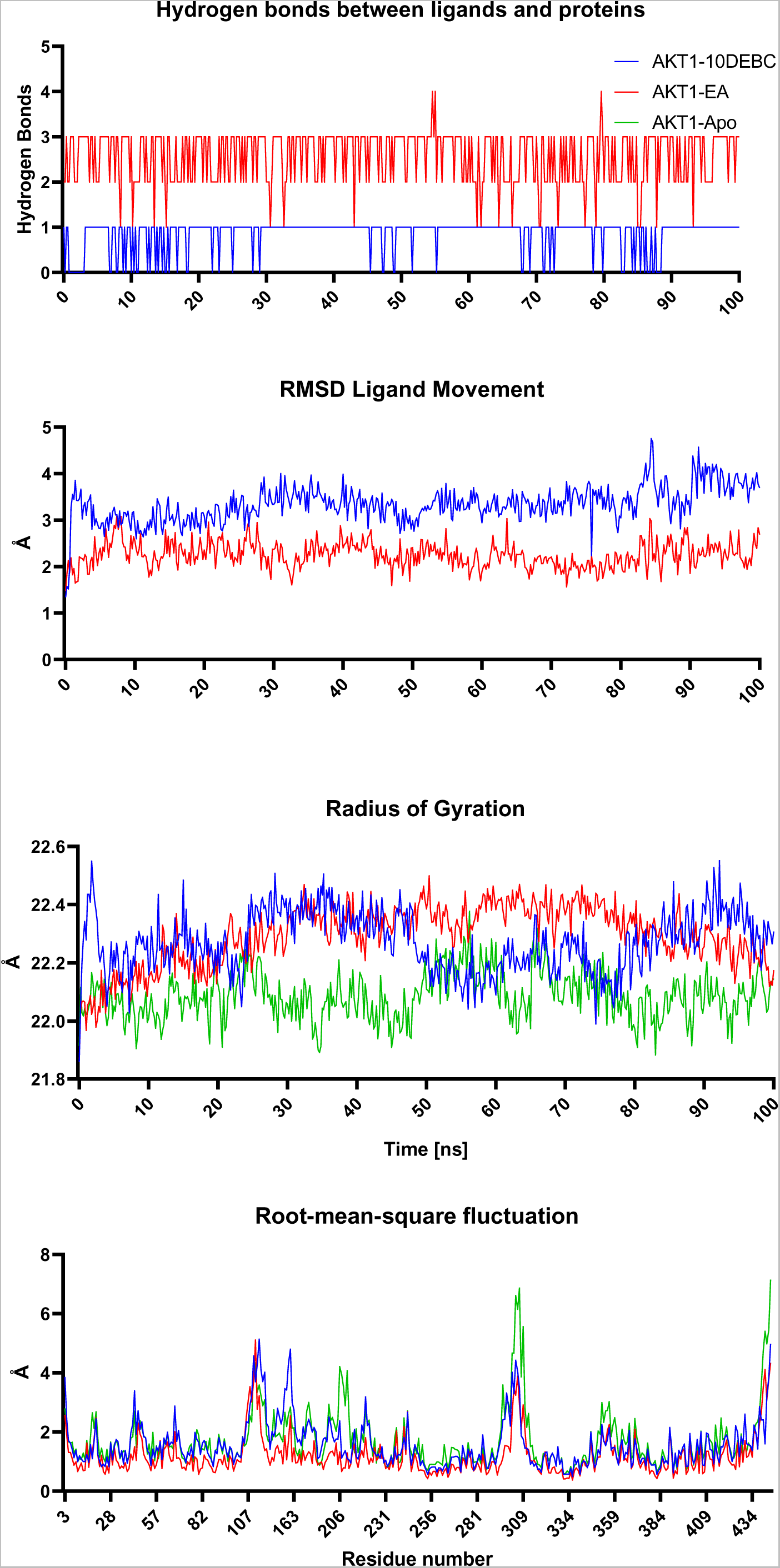
Molecular dynamics simulation analyses for AKT1 complexes. RMSD: root-mean-square distance.

**Figure S2.**
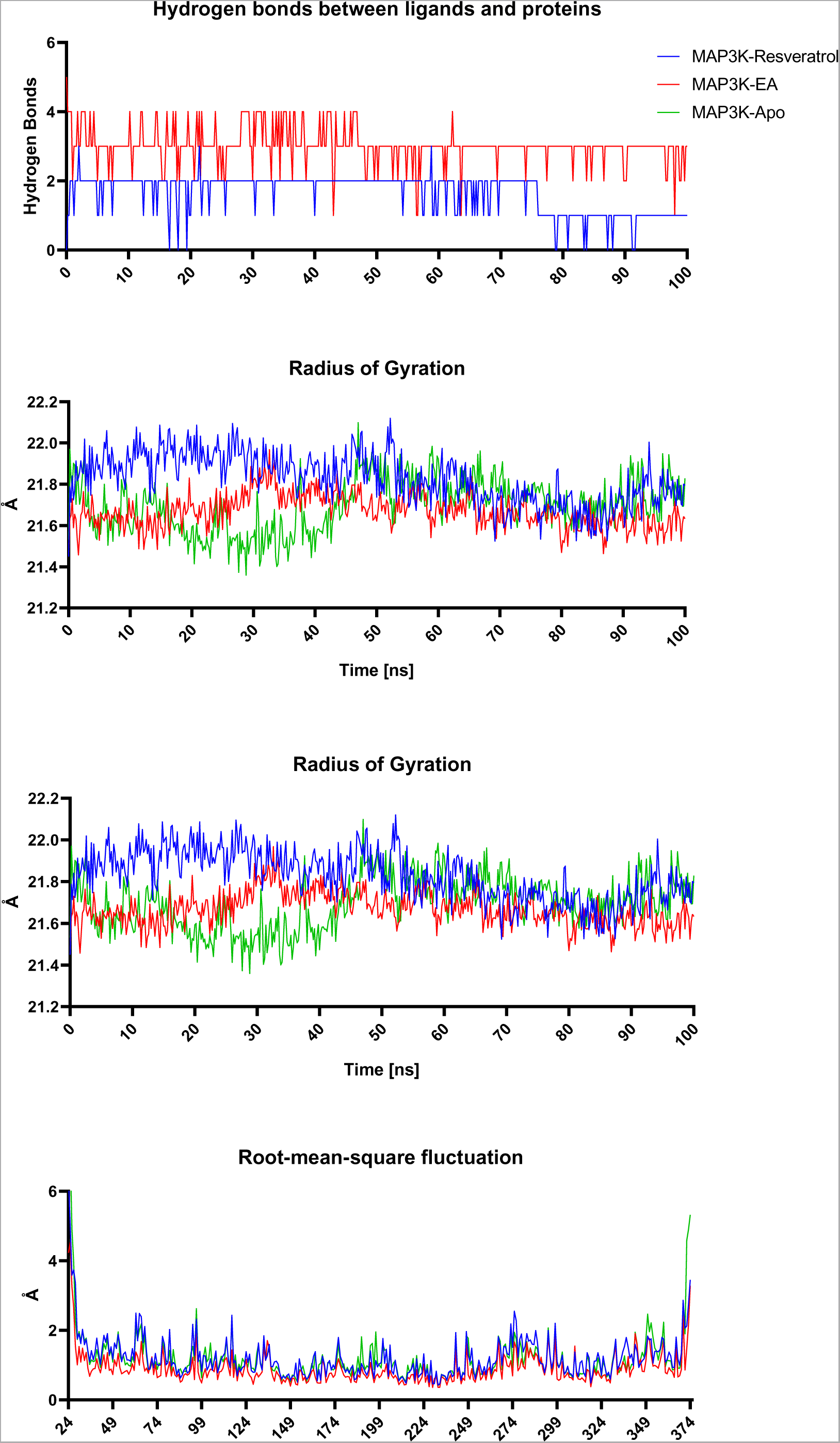
Molecular dynamics simulation analyses for MAP3K complexes. RMSD: root-mean-square distance

**Figure S3.**
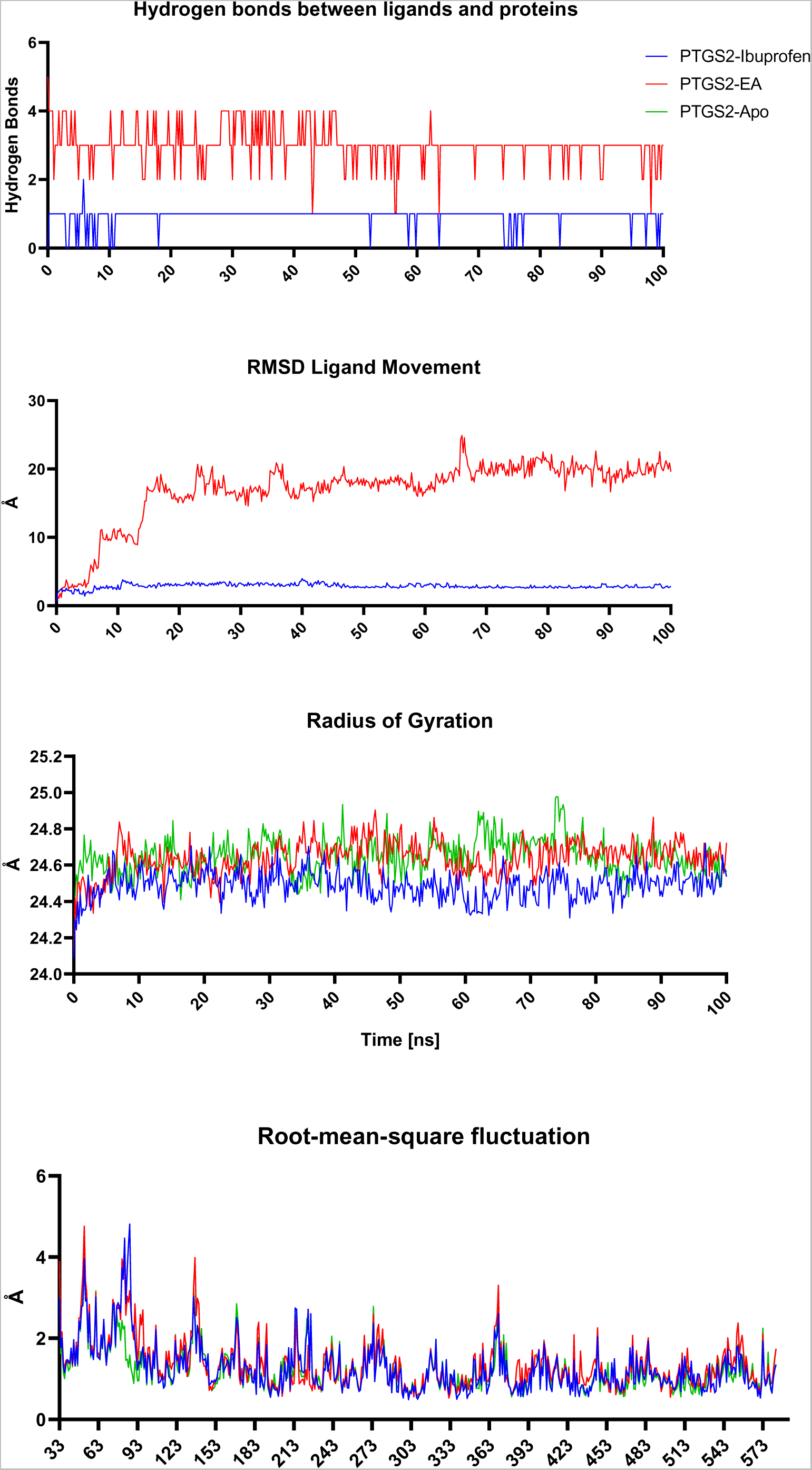
Molecular dynamics simulation analyses for PTGS2 complexes. RMSD: root-mean-square distance

**Figure S4.**
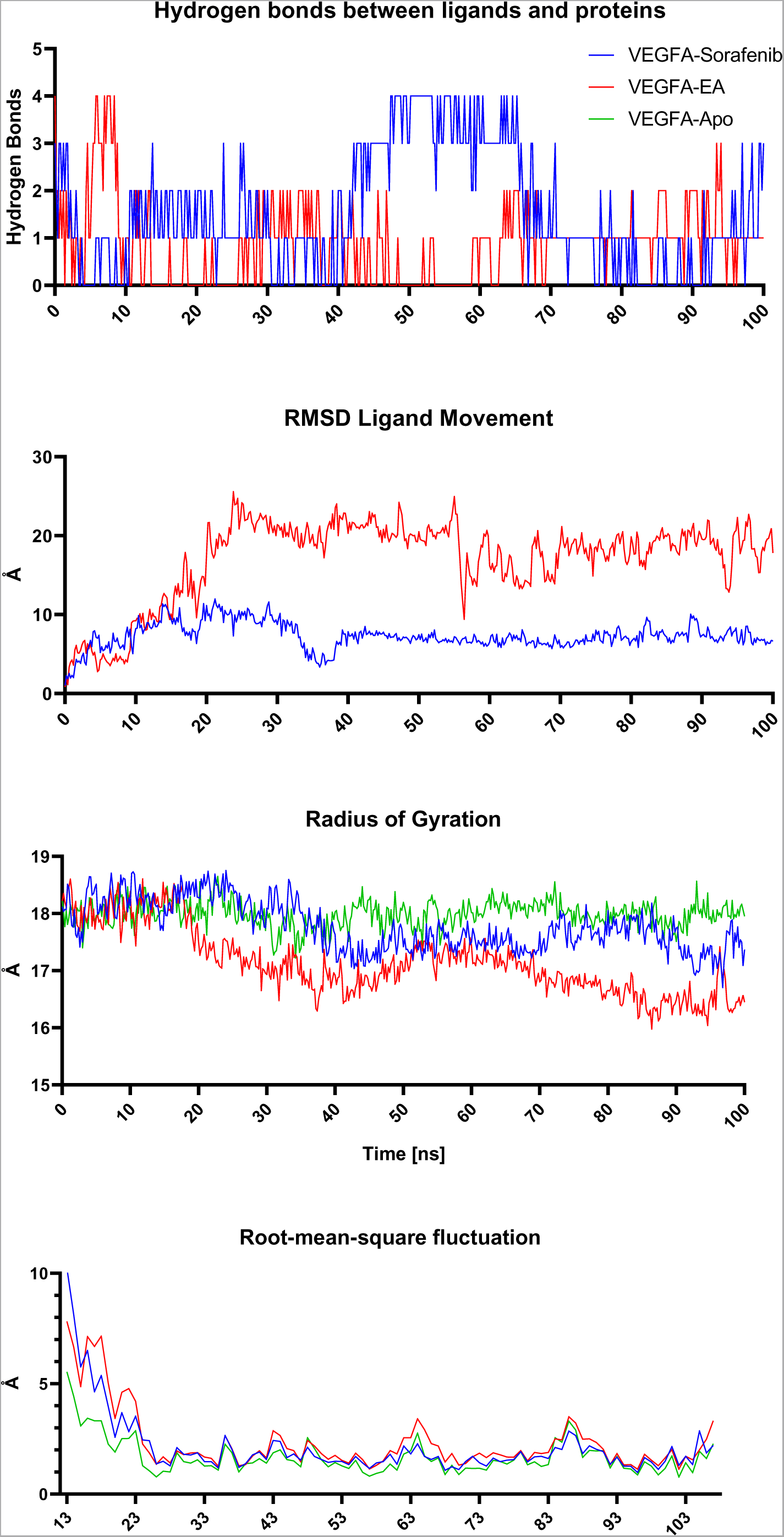
Molecular dynamics simulation analyses for VEGFA complexes. RMSD: root-mean-square distance.

**Figure S5.**
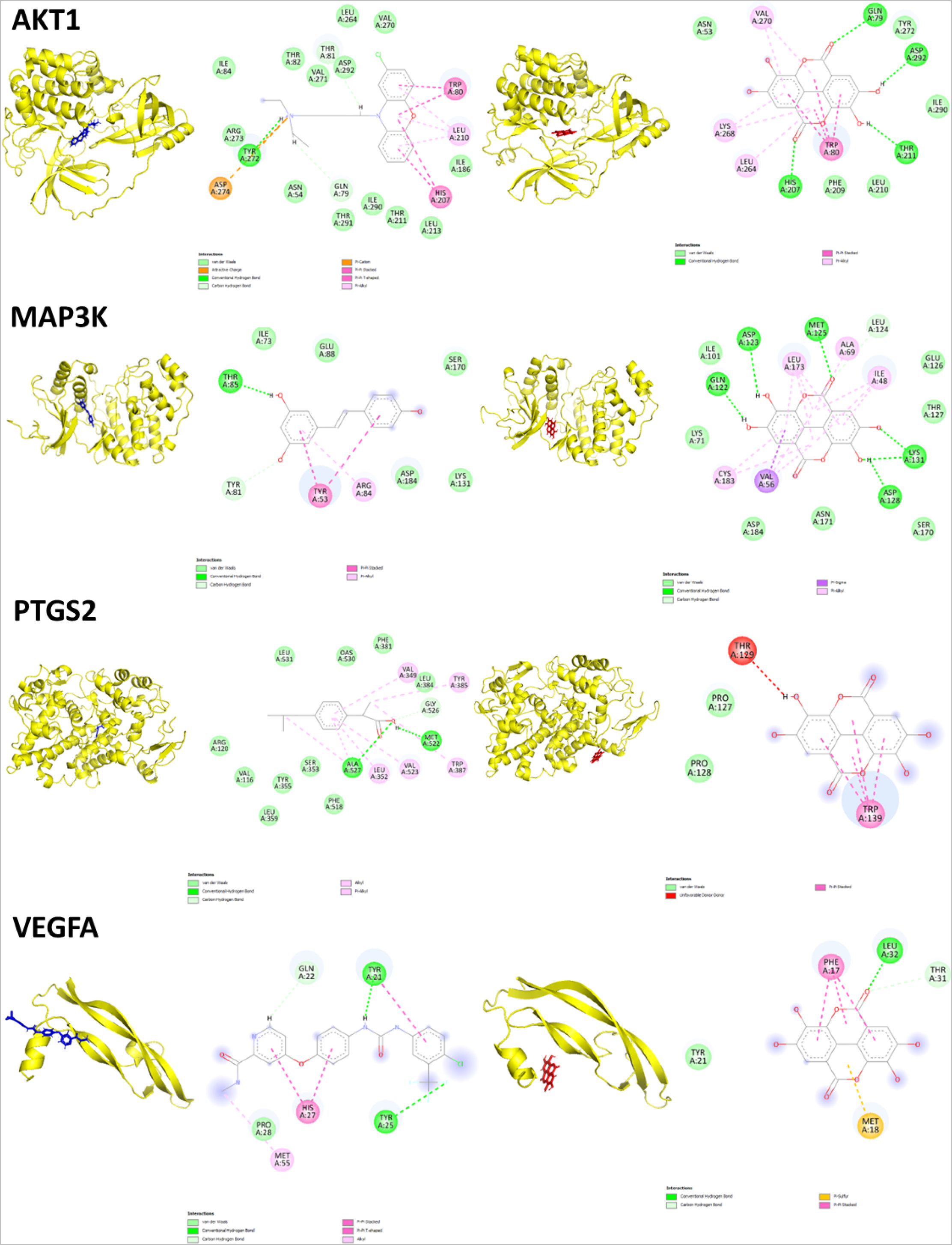
Binding modes of EA (red) and known binders (blue) towards their target proteins (yellow) at the 100^th^ ns of the simulations.

## Author contribution

SHH planned the study, performed bioinformatics analysis, and wrote the original draft. SHH, EML, and HD revised the manuscript and performed additional bioinformatics analysis. All authors have read the manuscript and agreed upon submission.

## Abbreviations

AKT1: AKT serine/threonine kinase 1
ALB: Albumin
DMNC: Density of Maximum Neighborhood Component
EA: Ellagic Acid
EPC: Edge Percolated Component
GO: Gene Ontology
HIF: Hypoxia-inducible Factor
ICAM: Intercellular Adhesion Molecule
IL: Interleukin
KEGG: Kyoto Encyclopedia of Genes and Genomes
MAPK: Mitogen-activated Protein Kinase
MCC: Maximal Clique Centrality
MMP: Matrix Metallopeptidase
MM-PBSA: Molecular Mechanics Poisson-Boltzmann Surface Area
MNC: Maximum Neighborhood Component
NF-κB: Nuclear factor kappa B
PPI: Protein Protein Interaction
PTGS: Prostaglandin-endoperoxide Synthase 2
SELP: Selectin P
TNF: (TNF-α) Tumor Necrosis Factor alpha
VEGFA: Vascular Endothelial Growth Factor A

